# High spatial resolution ^23^Na-MRI for ischemic brain injury detection

**DOI:** 10.64898/2026.02.05.703829

**Authors:** Yuanyuan Jiang, Xiaoqing Alice Zhou, Takahiko Imai, Lydia Hawley, David Hike, Sohail Mohammed, Grace Yu, Cenk Ayata, David Y. Chung, Xin Yu

## Abstract

High spatial resolution sodium (^23^Na) imaging of brain lesions remains challenging due to the intrinsically low signal-to-noise ratio (SNR) of ^23^Na-MRI compared with conventional proton (^1^H) MRI. In this study, we established a high-resolution ^23^Na-MRI platform based on 14 T preclinical scanner using a dual-tuned head-implanted RF coil. This configuration enables the acquisition of ^1^H-based T_2_-weighted anatomical and diffusion-weighted imaging (DWI), as well as ^23^Na-MRI, from the same animal. Brain-wide ^23^Na-MRI with 300×300 µm^2^ in-plane resolution was performed within the first 4 hours after acute middle cerebral artery occlusion (MCAO) induced ischemic stroke induction. Hyperintense ^23^Na-MRI signals were observed in the ischemic brain regions, presenting a contrast-to-noise ratio (CNR) that exceeded that of T_2_-weighted images. Infarct regions can be well-identified in the elevated ^23^Na signal, which is spatially consistent with regions of reduced apparent diffusion coefficient (ADC) values, indicating water restriction in the ischemic core. This work established implanted double-tuned ^1^H/^23^Na RF coils as a powerful approach for high-resolution ^23^Na-MRI readout, facilitating the early detection of ischemic brain injury.

## Introduction

Proton-based magnetic resonance imaging (^1^H-MRI) has provided critical insights into the spatiotemporal evolution of ischemic brain injury, with diffusion- and relaxation-based contrasts serving as sensitive indicators of cytotoxic edema and tissue damage^1-3^ . These contrasts, however, primarily reflect secondary changes in water mobility and tissue microstructure rather than the underlying failure of cellular homeostasis that defines irreversible injury. Loss of transmembrane sodium gradients is directly involved in cerebral ischemia. The resulting accumulation of sodium ions reflects impaired Na^+^/K^+^-ATPase activity, membrane depolarization, and breakdown of ionic compartmentalization, all of which are tightly linked to cellular viability^4^. Sodium MRI (^23^Na-MRI), therefore, provides a biophysically meaningful contrast by directly mapping tissue sodium distributions in vivo^5-11^. Despite this conceptual advantage, the interpretation of ^23^Na-MRI in ischemic stroke has been limited by insufficient spatial resolution and sensitivity.

The intrinsically low sensitivity of ^23^Na-MRI, approximately 20,000–30,000 times lower than that of ^1^H-RI^12,13^, severely constrains signal-to-noise ratio (SNR) and spatial resolution^14,15^. Consequently, prior studies in rodent stroke models have largely reported global or regional sodium signal changes, with limited ability to resolve heterogeneity within infarct cores or penumbral regions^16-20^. Even at ultra-high magnetic field strengths, e.g., 21 Tesla (T)^21,22^, voxel sizes remain substantially larger than those used in diffusion- or T_2_-weighted imaging, complicating direct biophysical comparisons between sodium accumulation and established proton-based markers of ischemic injury.

In this study, we tackled these limitations by combining ultra-high-field MRI at 14 T with a dual-tuned, head-implanted radiofrequency (RF) coil optimized for ^1^H/^23^Na imaging in a mouse model of middle cerebral artery occlusion (MCAO). This approach enables submillimeter-resolution ^23^Na-MRI, allowing spatially resolved comparisons between sodium accumulation, diffusion restriction, and T_2_ changes within ischemic tissue. By improving both sensitivity and spatial fidelity, our results establish a framework for interrogating sodium-based biophysical mechanisms of ischemic injury at a scale previously accessible only to proton-based MRI.

## Results

### Head-implanted RF coils enable high-spatial-resolution ^23^Na-MRI of the mouse brain

Building on our previous development of head-implanted RF coil technology^23,24^, we implanted a lightweight (<2.5 g), dual-tuned RF coil directly onto the mouse skull (**Fig. 1A**). The coil was able to be tuned to both ^1^H and ^23^Na resonance frequencies (**Fig. 1B**), enabling switching between proton- and sodium-based acquisitions on a 14 T preclinical MRI system. This configuration allowed B_0_ shimming and slice prescription to be optimized using high-SNR ^1^H-MRI and subsequently applied to ^23^Na-MRI acquisitions without geometric mismatch.

**Figure 1.**
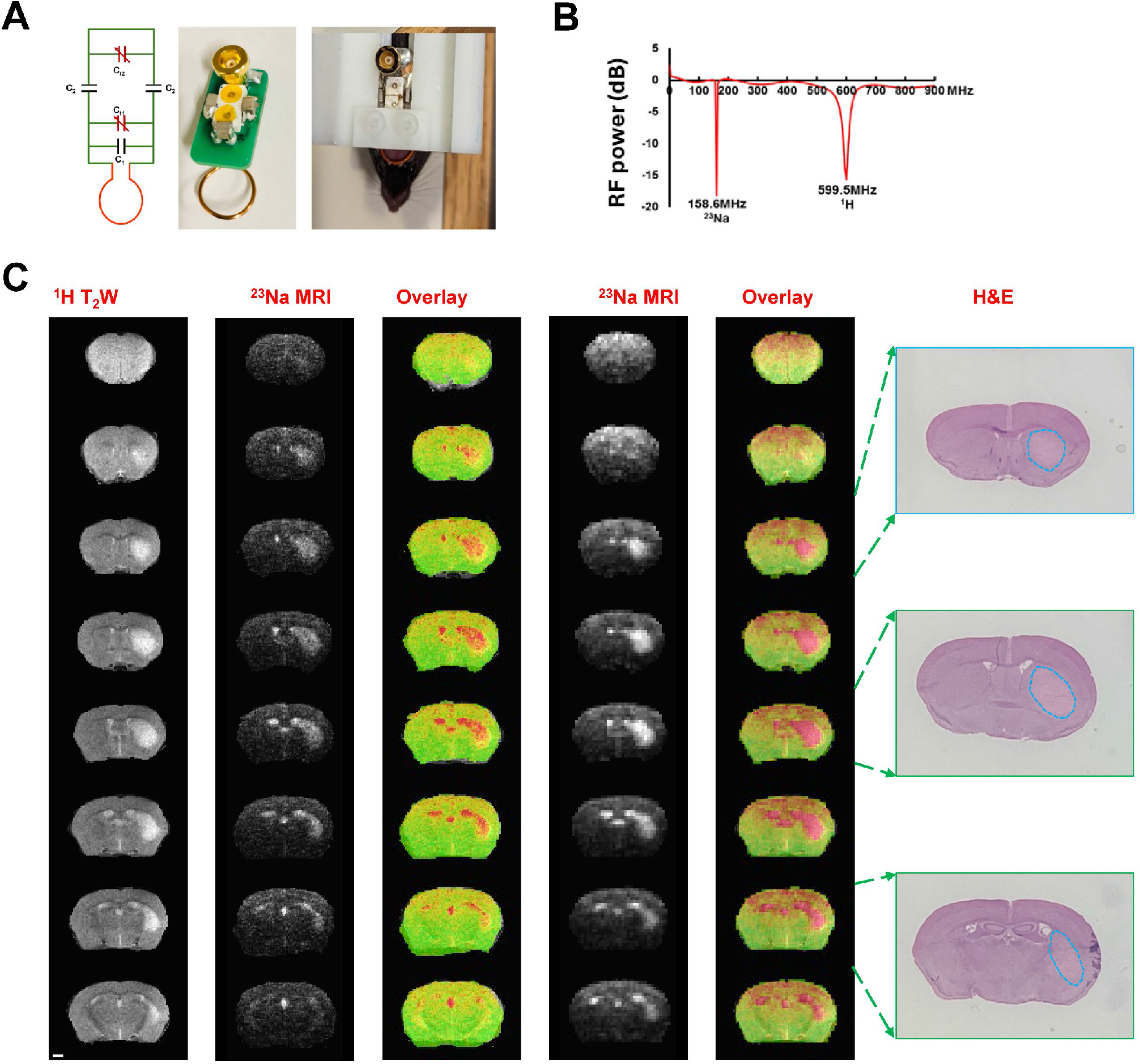
High-resolution ^1^H-MRI and ^23^Na-MRI for MCAO ischemic stroke mouse mapping with multimodal fMRI platform. (**A**) The dual-tuned RF coil was implanted into the mouse’s skull to acquire ^1^H and ^23^Na-MRI signals. (**B**) The dual-tuned RF frequency for both ^1^H and ^23^Na-MRI acquisition. (**C**) Specifically designed RF coils allow the acquisition of ^1^H T_2_-weighted (T_2_W) anatomical and ^23^Na-MRI with two spatial resolutions (150x150x500 µm^3^ for 115 min acquisition and 300×300×300 µm^3^ for 128 min acquisition) from the same MCAO ischemic stroke mouse in a 14T scanner. The brain lesion can be confirmed by H&E staining.

Using this approach, we achieved high-resolution ^23^Na-MRI in mice subjected to MCAO. Representative images acquired at spatial resolutions of 150 × 150 × 500 µm^3^ (115 min acquisition) and 300 × 300 × 300 µm^3^ (128 min acquisition) revealed clear sodium hyperintensity in ischemic regions 24 hours after stroke induction (**Fig. 1C**). These sodium-defined lesions spatially overlapped with infarct regions identified on T_2_-weighted ^1^H-MRI and were independently confirmed by hematoxylin and eosin (H&E) staining. Notably, the achieved sodium imaging resolution approached that of conventional proton-based anatomical imaging, enabling direct visual correspondence between modalities.

To evaluate the temporal evolution of sodium accumulation during the acute phase of ischemia, we performed serial ^23^Na-MRI in a representative MCAO mouse using a 300 × 300 µm^2^ in-plane resolution (**Fig. 2A**). Dynamic imaging revealed a progressive increase in ^23^Na signal intensity within the infarct region over time, with images acquired at 18.7-minute intervals following stroke induction (**Fig. 2B)**. Quantitative analysis demonstrated a ∼112% increase in ^23^Na signal within the infarct core over a 2.5-hour measurement period, while sodium signal in the contralateral hemisphere remained stable (**Fig. 2C**). These results demonstrate the feasibility of using high-resolution ^23^Na-MRI to map spatially and temporally evolving brain lesions with submillimeter precision during the early stages of ischemic stroke.

**Figure 2.**
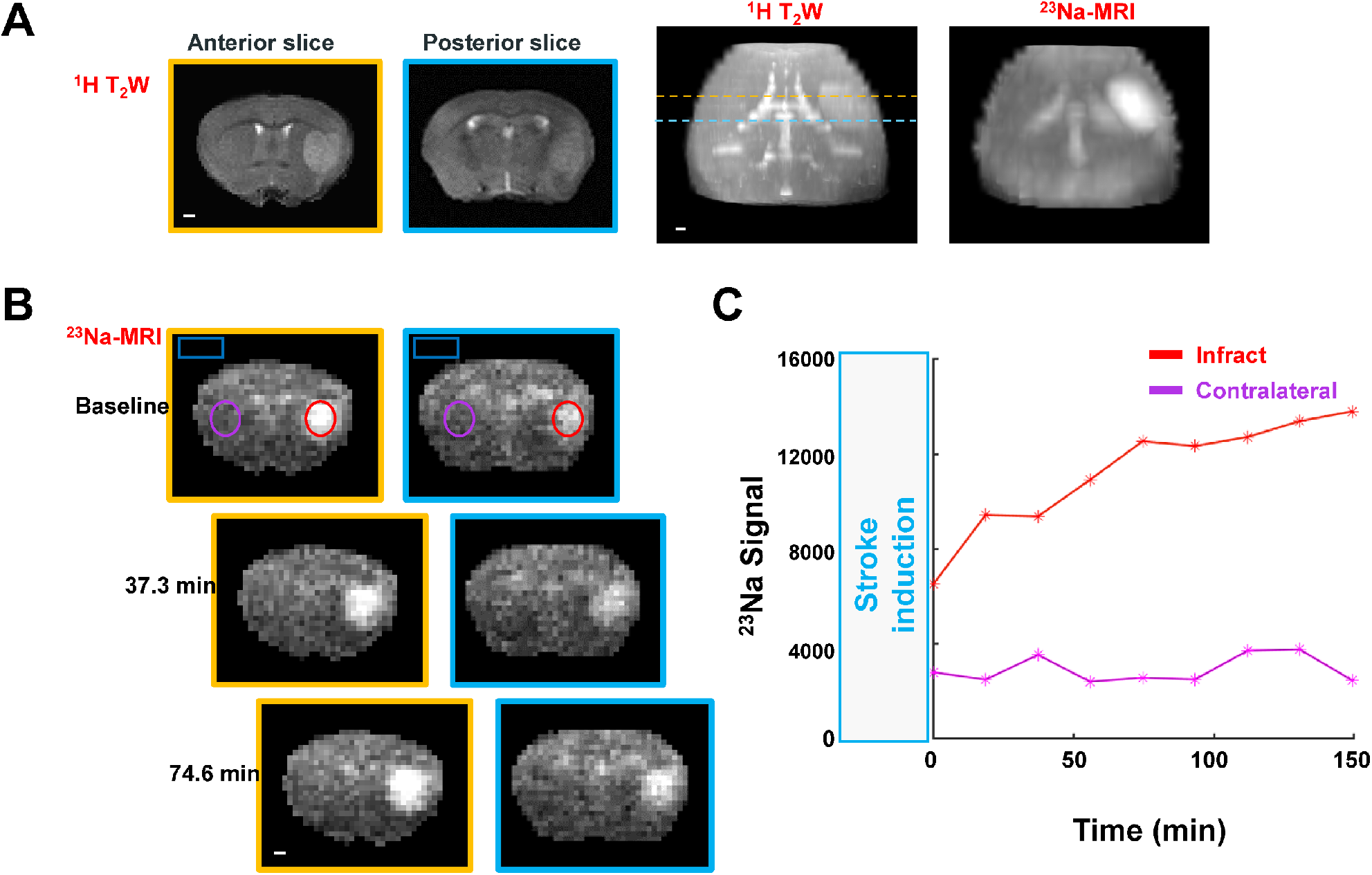
Temporal dynamics of ^23^Na-MRI in an acute MCAO study. (**A**) Representative coronal T_2_W slices following the transient MCAO stroke induction. The slices correspond to the selections in 3D volumetric T_2_W and ^23^Na-MRI images. **(B**) Representative ^23^Na-MRI slices at different time points after the ^23^Na-MRI measurement (3 hours after transient MCAO surgery). **(C**) The timecourse of ^23^Na signal from the infarct ROIs region and the contralateral hemisphere of the ^23^Na-MRI image (**B**). The ROIs were delineated from the hyperintense ^23^Na slices (300×300 µm^2^ in-plane resolution).

### Quantitative characterization of ^23^Na signal enhancement in ischemic lesions

To quantify lesion-associated sodium signal changes, we performed ^23^Na-MRI in mice within the first four hours following MCAO induction. Sodium images were acquired using a multi-slice 2D FLASH-based sequence with an in-plane resolution of 300 × 300 µm^2^ and a slice thickness of 500 µm. Marked hyperintensity was consistently observed in ischemic regions on ^23^Na-MRI (**Fig. 3A**). For comparison, T_2_-weighted ^1^H images were acquired using a RARE sequence with matched slice geometry (150 × 150 µm^2^ in-plane resolution, 500 µm slice thickness).

**Figure 3.**
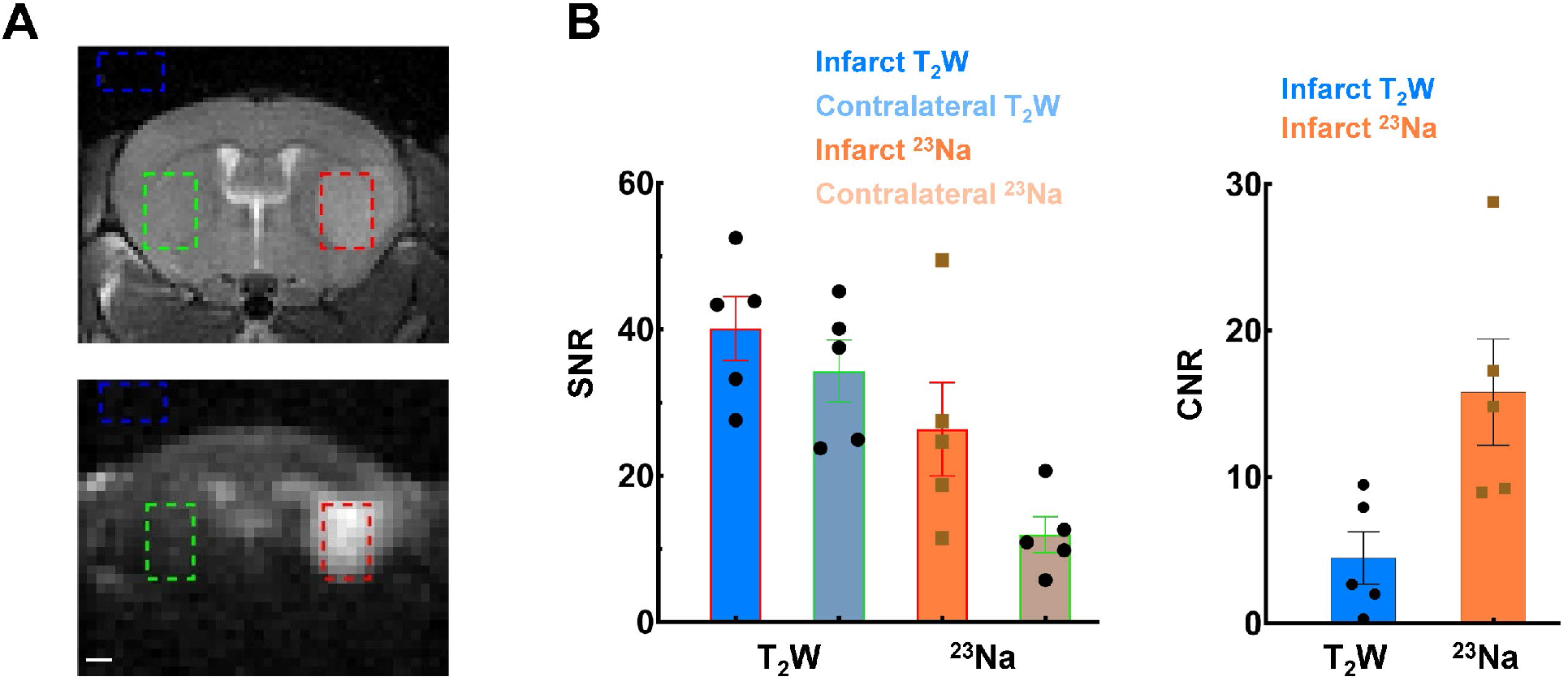
Quantitative measurements of the ^23^Na-enhanced signals in lesioned areas. (**A**) Following the stroke induction, hyperintense ^23^Na signals were identified in the lesioned brain areas compared to RARE images. The ROIs were selected from the lesioned area (red rectangle contour), the contralateral ROI (green), and the background outside the brain (blue) for SNR and CNR calculation. (**B**) The SNR of the ^23^Na-enhanced signals (26.34 ± 5.71) remained lower than the proton-based RARE images (40.10 ±3.91). There were no significant differences in SNR between the infarct and contralateral T_2_W images, while a significance was observed in the ^23^Na images (p=0.03, paired Wilcoxon t-test, n=5 mice, mean ± SEM). The CNR from hyperintense ROIs (15.80 ± 3.20) was more than three times higher than the T_2_W images (4.47 ± 1.59) from the infarct region (p=0.04, n=5 mice, mean ± SEM). The scale bars (1 mm) are denoted in the figure.

Although the absolute SNR of ^23^Na-MRI in lesion regions (26.34 ± 5.71) was lower than that of proton-based RARE images (40.10 ± 3.91), the contrast-to-noise ratio (CNR) between infarct and contralateral regions was substantially higher for ^23^Na-MRI (15.80 ± 3.20) compared with T_2_-weighted imaging (4.47 ± 1.59; **Fig. 3B**). These findings indicate that, despite lower intrinsic SNR, ^23^Na-MRI provides superior lesion contrast sensitivity relative to conventional proton-based T_2_-weighted imaging.

### Comparison of ^23^Na-MRI with diffusion-based ADC mapping

To further validate the lesion specificity of sodium-based contrast, we compared ^23^Na-MRI with diffusion-derived apparent diffusion coefficient (ADC) maps in the same animals. Both modalities consistently identified ischemic regions across multiple slices, with lesion locations corroborated by TTC staining (**Fig. 4A**). Across all animals, the injured hemisphere exhibited a pronounced increase in ^23^Na signal intensity relative to the contralateral side. Spatial correspondence between elevated sodium signal and reduced ADC values was observed within infarct cores (**Fig. 4B**), indicating overlap between sodium accumulation and diffusion restriction. However, group analysis of line profiles revealed that ^23^Na-MRI produced a more spatially confined lesion profile with much better CNR compared with ADC maps (**Fig. 4C**). Additionally, three-dimensional volumetric renderings further illustrated the enhanced lesion delineation provided by ^23^Na-MRI relative to diffusion imaging (**Supplementary Fig. 1**). Together, these results demonstrate that high-resolution ^23^Na-MRI offers robust sensitivity and spatial specificity for characterizing cortical and subcortical brain lesions following ischemic stroke.

**Figure 4.**
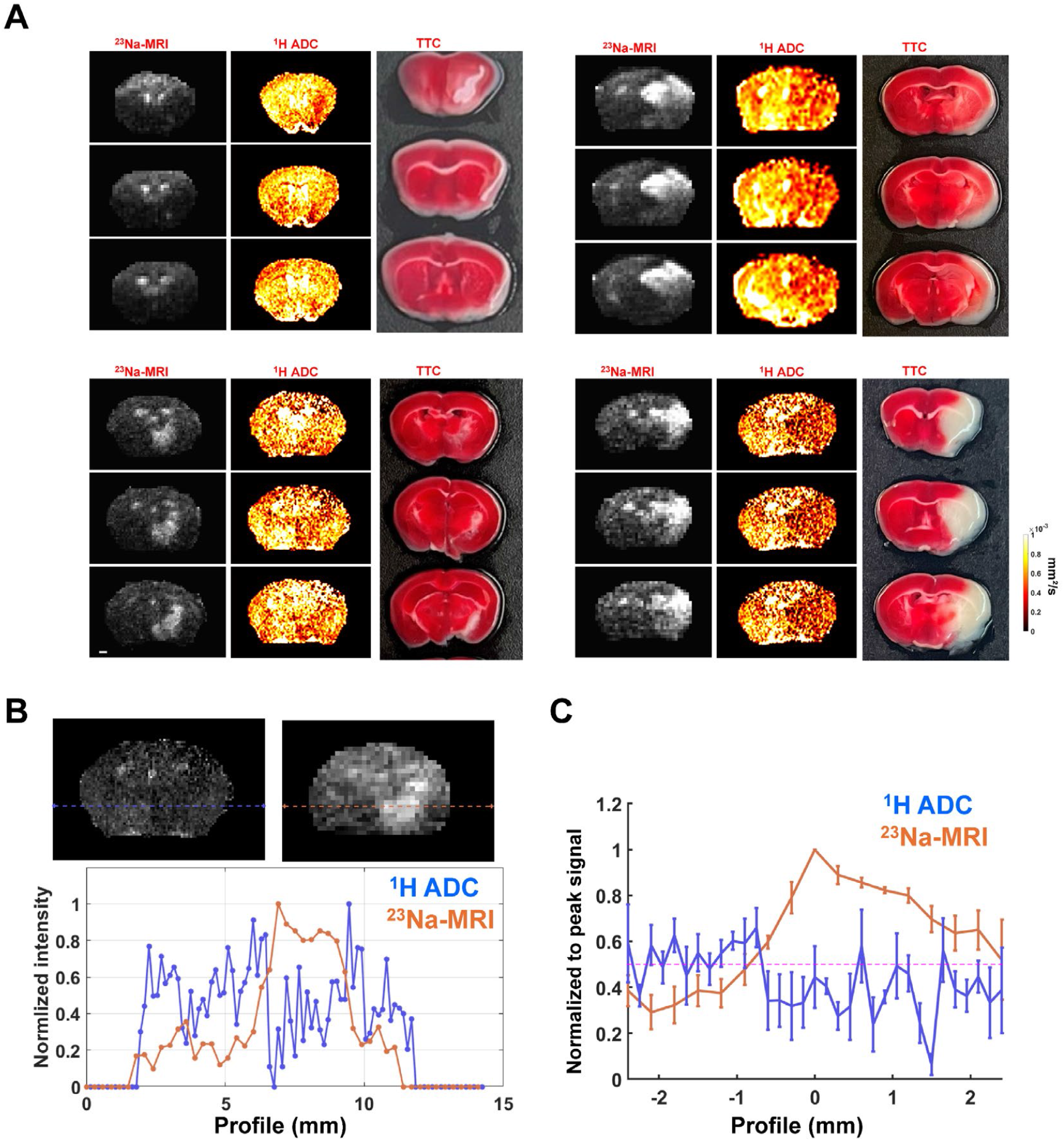
Comparison of ADC mapping and ^23^Na-MRI in brain injuries. (**A**) Brain lesions can be well-identified in the ^23^Na image in the subcortical regions, which could also be detectable in diffusion-based ADC maps, although variations in T_2_-weighted maps and TTC staining. (**B**) Representative signal changes across the infarct regions are shown in the ADC map and ^23^Na image, normalized to peak values. (**C**) The spatial distribution of ^23^Na and ADC values, aligned based on the peak ^23^Na-MRI signal from the infarct core regions (mean ± SEM, n = 4 mice). Scale bars, indicating 1 mm, are denoted within the figures.

## Methods and Data Analysis

### Animals

The C57BL/6 mice (Charles River Laboratories, MA) were housed (group 3-4/cage) under a 12-h light/dark cycle with food and water ad libitum. They were maintained under a 12-hour light/dark cycle with unrestricted access to food and water. All animal procedures were performed in compliance with protocols approved by the Animal Care and Use Committee (IACUC) at Massachusetts General Hospital (MGH), and the animals were cared in accordance with the National Research Council’s Guide for the Care and Use of Laboratory Animals.

### Experimental workflow

The RF coil was head-implanted one week before the MCAO stroke surgery induction. After the stroke induction, the mice were maintained on the warming pad for 15 minutes. Then, the mice were positioned on a support 3D-printed cradle for MRI scanning.

### RF implantation

The general procedure for RF implantation is described previously^24^. Briefly, mice were anesthetized with isoflurane 2.5% for induction and 1%–1.25% for maintenance (70% N_2_/30% O_2_). The mouse head was fixed in a stereotaxic device, and the craniotomy incision (the size of an RF ring) was exposed to position the coil on the skull. The cyanoacrylate glue and dental cement (Stoelting Co., Wood Dale, IL) were applied to secure the coil and to firmly cover the underlying brain.

### The focal cerebral ischemia stroke model

C57BL/6 male mice were subjected to either permanent or 60-minute transient MCAO (pMCAO or tMCAO). Surgery was performed during the inactive phase/light cycle (7:00 to 19:00). General surgical preparation was performed as described previously^25,26^. MCAO was performed under isoflurane anesthesia (2.5-3% induction, 1.5-2% maintenance in 70% N_2_O/30% O_2_), and body temperature was maintained at 37±0.5°C using a heating pad. A laser Doppler flowmetry (LDF) probe was attached to the right middle cerebral artery (MCA) territory (2 mm posterior, 2 mm lateral from the bregma) to monitor cerebral blood flow (CBF). The ventral midline neck was incised, and the right common carotid artery (CCA) bifurcation was exposed without damaging the nerves. After ligation of CCA, the external carotid artery (ECA) was cut at the middle of both ligations and then a nylon monofilament (602212PK10Re, Doccol Corporation, Sharon, MA, USA) was inserted into the ECA and advanced to the MCA origin through the internal carotid artery (ICA). In the tMCAO group, the CCA was opened after monitoring CBF persistence, and then the mouse was kept under anesthesia during occlusion periods. At 60 min after tMCAO, the filament was removed to restore CBF as reperfusion. On the other hand, the CCA was kept closed in the pMCAO group and not moved to the filament until sacrifice. Immediately after surgery, saline was IP administered at 10 ml/kg to prevent dehydration. The mice were then transferred to a warm recovery chamber and scanned by MRI for 4 hours after surgery. After scanning, mice were then sacrificed to analyze infarct volume. In total, 7 mice were used and divided; 3 mice were tMCAO, and 4 were pMCAO.

### ^1^H-MRI and ^23^Na-MRI acquisiton

All images were acquired with a 14 T horizontal bore magnet (Magnex Sci, UK), interfaced to a Bruker AV-Neo console and ParaVision 360 V.3.3 (Bruker, BioSpin GmbH, Germany), and equipped with a 6 cm microimaging gradient set, capable of providing 1.2 T/m (Resonance Research, MA). A ∼6 mm dual-tuned transceiver surface implantable coil (MRIBOT LLC, MA) was used to acquire MRI images. The dual-tuned surface coil was tuned and matched outside of the magnet bore. The ^1^H-MRI scans were acquired to verify the correct animal positioning and shimming inside the magnet. The T2-weighted (T_2_W) RARE and diffusion-weighted imaging (DWI) scans were acquired first. Magnetic field B_0_ homogeneity was optimized first by automatic shimming from ^1^H acquisition and then used for ^23^Na shimming. For the subsequent ^23^Na-MRI scan, the optimized RF power for the ^23^Na-MRI scans was measured using the AdjRefPowX report provided by the Bruker sequence.

The T_2_W anatomical, DWI, and ^23^Na-MRI scans were acquired with the same FOV (14.4×9.6×8 mm^3^, slice thickness=0.5 mm) unless otherwise indicated. A modified ultrafast FLASH-based fMRI 2D line-scanning pulse sequence was applied for ^23^Na-MRI acquisition^27^. The pulse sequence scripts and binary code implementations of the 2D line-scanning method are available at our shared source: https://tnnc.mgh.harvard.edu/method. The parameters for ^23^Na-MRI are applied following: TE=1.2 ms, TR=10 ms, matrix= 48×32×16, resolution=300×300×500 µm^3^. For T_2_W, a RARE sequence was implemented to acquire anatomical images with the same geometry. TE=12 ms, TR=1500 ms, RARE factor=6, averages=16, matrix =96×64×16, resolution=150×150×500 µm^3^. For DWI, TE=14 ms, TR=2500 ms, b values of 0, 50, 100, 150, 250, 400, and 600 s/mm^2^ were applied and 96×64×16, resolution=150×150×500 µm^3^. During the scanning, the mice were anesthetized with isoflurane 1%–1.25%. The body temperature was maintained at 37^°^C by blowing warm air through the bore and recording with a rectal probe (Small Animal Monitoring Model 1030, SA Instruments, Inc., NY, USA).

### Brain coronal section staining for ischemic area

To evaluate the tissue outcome, mice brains were cut at coronal section and stained by 2,3,5-triphenyl-tetrazolium chloride (TTC) or Hematoxylin and Eosin (H&E).

### TTC staining

Mice were euthanized by cervical dislocation under deep anesthesia (5% isoflurane). Brains were carefully removed and sectioned into 1 mm-thick coronal slices and immersed in 2% of 2,3,5-triphenyl-tetrazolium chloride (TTC; Sigma-Aldrich, Inc., St. Louis, MO, USA) saline until intact tissue was sufficiently stained.

### H&E staining

Mice were deep anesthetized with 5% isoflurane and systemically perfused with saline for 2 minutes via the left ventricle. Brains were carefully removed, frozen in isopentane on dry ice, and stored at -80 degrees. Twenty micrometer-thick cryosections were collected every 1 cm on ionized glass slides and stained with hematoxylin and eosin (H&E; Poly Scientific R&D Corp., Bay Shore, NY, USA).

### SNR and CNR measurements

The SNR was computed by averaging the signal from the hyperintense voxel delineating the center of the ischemic lesion regions and dividing by the standard deviation of the voxel from the background outside the brain region (**Fig. 2A & Fig. 3A**). The contralateral ROI voxels were positioned on the corresponding contralateral normal area with the same ROI size. The CNR was computed from the hyperintense ^23^Na area and contralateral ROIs, divided by the standard deviation of the background region^28^. For SNR and CNR calculation, the RARE slice was downsampled to the same resolution as the ^23^Na-MRI map. The ADC maps were generated from the nonlinear least-squares fit through all b values for each slice acquired by DWI. The data processing and analysis methods were performed with Analysis of Functional NeuroImages (AFNI) (NIH, Bethesda, USA) and MATLAB (MathWorks, Natick, USA). The contralateral ratio was determined by the hyperintense ^23^Na infarct area and corresponding contralateral ROIs from the acquired RARE, ADC, and ^23^Na images. For the group analysis, a Wilcoxon t-test was performed. The error bars in all figures represent the mean ± standard error of the mean (SEM) unless otherwise indicated. The threshold for statistical significance was set at p < 0.05.

## Discussion

Using ultra–high-field MRI in combination with a head-implanted, dual-tuned RF coil, we demonstrate submillimeter-resolution ^23^Na-MRI of the mouse brain *in vivo* and apply this capability to characterize ischemic injury following MCAO. This technical advance enables robust visualization of spatially confined sodium accumulation within infarcted tissue, presenting more salient lesion contrast when compared with conventional proton-based T_2_-weighted imaging or diffusion-derived ADC maps. Together, these findings establish high-resolution ^23^Na-MRI as a sensitive and biophysically meaningful complement to standard ^1^H-MRI for preclinical stroke research.

### Overcoming the intrinsic SNR limitations of ^23^Na-MRI

The principal obstacle to *in vivo* ^23^Na-MRI is its intrinsically low SNR, which arises from multiple fundamental factors, including the lower gyromagnetic ratio of ^23^Na relative to ^1^H, its substantially lower tissue concentration compared with water protons, and quadrupolar relaxation that produces rapid biexponential T_2_/T_2_* decay^5,29-31^. Together, these constraints have historically necessitated large voxel sizes and long acquisition times, limiting spatial resolution and obscuring fine-scale heterogeneity within ischemic lesions. In the present study, we addressed these limitations by combining ultra-high magnetic field strength (14 T) with a skull-mounted implantable RF coil optimized for dual-nucleus (^1^H/^23^Na) operation. The 14 T enables increased SNR based on the supra-linear scaling factor with field strengths (SNR ∼ B_0_^1.65^ for coil arrays and B_0_^2.0^ for small surface coils )^32,33^. And, the implantable coil configuration provides several key advantages: (i) reduced sensitivity to air–tissue susceptibility interfaces, improving B_0_ homogeneity and shimming performance at high field^34^; (ii) substantial SNR gain in superficial cortical regions due to the proximity of the receive element; and (iii) seamless switching between proton and sodium imaging with identical geometry, enabling precise multimodal co-registration. Collectively, these features allowed us to achieve in-plane resolutions down to 300 μm for ^23^Na-MRI—an order-of-magnitude improvement over typical preclinical sodium imaging protocols, even with 21 T scanners^16-20^. Nevertheless, implantable surface coils inherently exhibit limited penetration depth and B_1_ inhomogeneity, leading to signal attenuation in deeper brain structures. Despite these constraints, we demonstrate reliable detection of subcortical ischemic lesions, underscoring the feasibility of this approach for whole-brain sodium mapping in mice. Future RF sensing developments, including figure 8 coil geometries^24,35,36^, multi-element cryogenic arrays^37-39^, and wireless implantable RF coils with built-in varactor-based preamplifier^40,41^, could further extend sensitivity and depth coverage, enabling even finer spatial resolution and improved quantitative accuracy.

### High-resolution ^23^Na-MRI provides superior lesion contrast

A key observation of this work is that high-resolution ^23^Na-MRI yields markedly higher lesion-based CNR than conventional T_2_-weighted ^1^H-MRI, despite its lower absolute SNR. In human studies, the ^23^Na elevation is reported to be ∼50% above the contralateral hemisphere value^8,14,42,43^, whereas animal models show ∼100%^44^. By improving SNR, we were able to accurately delineate the ischemic injury volume (**Fig. 1 & Supplementary Fig. 1**) and detect a time-dependent ∼112% elevation of ^23^Na concentration within the infarct area with enhanced sensitivity (**Fig. 2C**). Also, our dynamic measurements demonstrate a gradual, time-dependent increase in ^23^Na signal within the infarct core during the acute post-ischemic period, highlighting the potential of sodium MRI to track lesion evolution rather than simply providing a static snapshot of tissue damage. This enhanced CNR reflects the strong biophysical coupling between sodium accumulation and the loss of cellular ionic homeostasis that defines irreversible ischemic injury (**Fig. 3**). By improving spatial resolution and minimizing partial-volume effects, we were able to delineate infarct boundaries with greater precision and detect heterogeneous sodium elevations within both cortical and subcortical regions.

### Relationship between sodium accumulation and diffusion restriction

Comparison with diffusion-derived ADC maps revealed substantial spatial overlap between regions of elevated sodium signal and reduced ADC, consistent with prior observations that both contrasts reflect ischemia-induced cellular dysfunction^45-48^. However, line-profile and volumetric analyses showed that ^23^Na-MRI produced more spatially lineated lesion profiles with higher CNR than ADC maps. This distinction likely reflects differences in the underlying biophysical mechanisms, that ADC reductions occur rapidly after ischemia onset due to cytotoxic edema and water redistribution^49,50^, whereas sodium accumulation reflects a slower breakdown of membrane integrity, Na^+^/K^+^-ATPase failure, and progressive loss of ionic gradients^44^. These differing temporal dynamics suggest that combined ADC and ^23^Na-MRI measurements could provide complementary information about tissue states. In principle, regions exhibiting diffusion restriction without substantial sodium elevation may correspond to metabolically compromised but potentially salvageable penumbral tissue, whereas areas with pronounced sodium accumulation likely represent irreversibly damaged infarct core (**Fig. 4**). Comparable to the conventional diffusion/perfusion weighted (DWI–PWI) mismatch framework^51-54^, a mismatch between ADC-defined lesions and ^23^Na-defined sodium accumulation could therefore offer an alternative strategy for identifying penumbral regions and estimating ischemic progression^14,18,42,43^.

### Biological interpretation and limitations

While an elevated ^23^Na signal is closely linked to loss of cellular viability, it is important to recognize that total sodium MRI does not distinguish intracellular from extracellular sodium pools. Multiple processes may contribute to sodium signal changes in ischemic tissue, including intracellular sodium influx, extracellular volume expansion due to vasogenic edema^42,55,56^, altered paravascular fluid transport^57^, and regional perfusion heterogeneity^6,58^. Disentangling these contributions will be essential for fully exploiting sodium MRI as a quantitative biomarker of tissue viability. Advanced methodological approaches, such as multiple-quantum filtering^59,60^, and optimized radial or ultrashort-echo trajectories^61-63^, offer potential avenues to increase sensitivity to intracellular sodium and to better separate compartmental contributions. Integration of these techniques with the dual-tuned RF platform presented here could substantially enhance the specificity of sodium-based imaging biomarkers.

## Conclusion

The present study demonstrates that technical barriers that have long limited ^23^Na-MRI can be overcome in preclinical settings, enabling spatial resolutions approaching those of conventional proton MRI. By directly probing ionic dysregulation, high-resolution sodium imaging provides information that is fundamentally distinct from hemodynamic, diffusion-based, or relaxation-based contrasts. In this study, T_2_-weighted imaging is interpreted as reflecting edema and late-stage infarction, whereas ADC/DWI is used to identify acute to subacute ischemic infarction. In contrast, ^23^Na-MRI provides complementary information that is not fully captured by conventional water-related structural and diffusion changes. When combined with diffusion and perfusion imaging, sodium MRI has the potential to refine the assessment of the infarct core and penumbra, improve estimates of stroke onset and progression, and ultimately enhance the translational relevance of preclinical stroke models.

## Acknowledgments

This research was funded by NIH funding (RF1NS113278, RF1NS124778, R01NS122904, R01NS120594), S10 instrument grant (S10 MH124733, S10OD036211), and the NSF grant 2123971 to Dr. Yu in Martino’s Center. Funding was received by the National Institute of Neurological Disorders and Stroke at the National Institutes of Health R01NS102969 to Dr. Ayata; R25NS065743, KL2TR002542, and K08NS11260 to Dr. Chung, Ellison Foundation for Dr. Ayata, Andrew David Heitman Foundation for Drs Ayata, Chung, and Jiang, American Heart Association and American Stroke Association (18POST34030369), Aneurysm and AVM Foundation Brain Aneurysm Foundation to Dr. Chung.

## Author contributions

Yuanyuan Jiang, Data curation, Software, Formal analysis, Validation, Investigation, Methodology, Funding acquisition, Writing–original draft; Xiaoqing Alice Zhou, Data curation, Formal analysis, Validation, Investigation, Methodology; Takahiko Imai, Data curation, Formal analysis, Validation, Methodology, Writing – original draft; Lydia Hawley, Methodology; David Hike, Methodology; Sohail Mohammed, Sohail Mohammed; Grace Yu, Methodology; Cenk Ayata, Resources, Data curation, Supervision, Funding acquisition; David Chung, Resources, Data curation, Supervision, Funding acquisition, Investigation, Visualization, Writing – review and editing; Xin Yu, Conceptualization, Resources, Data curation, Supervision, Funding acquisition, Validation, Investigation, Visualization, Project administration, Writing – review and editing.

