## Supplementary Figure 1 for "High spatial resolution ^23^Na-MRI for ischemic brain injury detection"

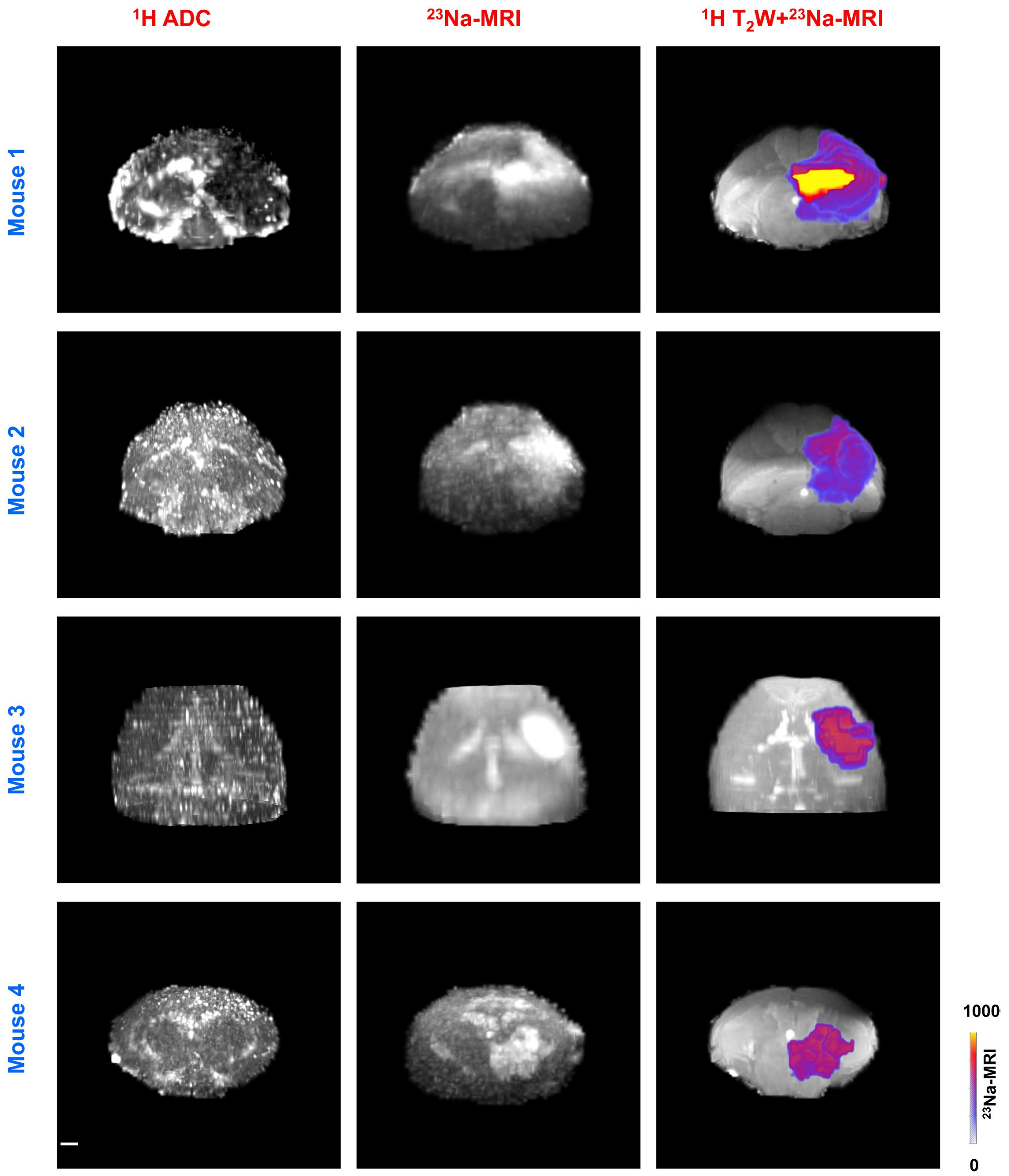


**Supplementary Figure 1.** The 3D volumetric display of ADC, $^{23}$Na-MRI mapping, and the hyperintense $^{23}$Na overlaid into the $^{1}$H based $T_2$W volume from 4 mice. Scale bars indicate 1mm and are denoted within the figures.
